# Physiological dominance of the scion in shaping root architecture under suboptimal temperature

**DOI:** 10.64898/2026.02.11.705274

**Authors:** Amnon Cochavi, Elad Oren, Fabian Baumkoler, Evgeny Smirnov, Ran Nisim Lati

## Abstract

**Background:** Non-optimal temperatures have become a major constraint on plant development under rapidly changing climatic conditions. Both sub- and supra-optimal temperatures reduce physiological activity, alter plant morphology, lead to plant mortality, and ultimately decrease crop productivity. Temperature-tolerant plants employ physiological and morphological mechanisms to mitigate such stress. In this study, we aimed to identify the source of temperature tolerance in warm-climate adapted melon (*Cucumis melo* L.).

**Results:** Suboptimal temperature–tolerant accession (Ananas Yoqne’am; AY) and susceptible accession (PI414723) were reciprocally grafted and grown under controlled temperature regimes of 16 °C, 25 °C, and 35 °C. Physiological and morphological traits were measured to characterize tolerance mechanisms and whole-plant responses. Temperature emerged as the dominant factor governing plant performance. Whereas non-grafted parental lines maintained consistent differences across all temperature regimes, reciprocal graft combinations diverged mainly under suboptimal (16 °C) conditions. Under these temperatures, scion identity strongly determined whole-plant performance through biochemical limitations.

**Conclusion:** These results highlight the importance of scion-derived traits in abiotic stress tolerance and their downstream influence on root function.

## 1. Introduction

Global climate change is intensifying the frequency, duration, and severity of temperature fluctuations, exposing agricultural systems to increasingly variable thermal environments. Temperature is among the most influential environmental factors governing plant physiology, exerting direct effects on photosynthesis, respiration, membrane stability, and metabolic processes (Bita and Gertas 2013). Sustained high temperatures may transiently enhance photosynthetic activity but commonly lead to oxidative stress, cellular injury, and premature senescence (Zhao et al. 2011; De la Haba et al. 2014). In contrast, low temperatures impose immediate constraints on physiological function, particularly by reducing stomatal conductance and carbon assimilation (Wilkinson et al. 2001; Theocharis et al. 2012). Such limitations, coupled with continued light absorption, promote the accumulation of reactive oxygen species (ROS), resulting in impaired membrane integrity, altered enzymatic activity, and accelerated foliar senescence (Suzuki and Mittler 2006; Savitch et al. 2009). Temperature stress also disrupts developmental trajectories, modifying root system architecture and above–belowground biomass allocation (Mokany et al. 2006; Nagel et al. 2009), ultimately constraining plant establishment and productivity.

Within this broader context, warm-season crops such as melon (*Cucumis melo* L.) are particularly susceptible to early-season thermal instability, especially during late-winter and early-spring sowing when rapid transitions between diurnal warmth and nocturnal cold are common. Although melon possesses substantial genetic and phenotypic diversity due to its domestication across a wide range of agroecological environments (Sebastian et al. 2010; Endl et al. 2018; Oren et al. 2022), the species remains highly sensitive to suboptimal temperatures during early developmental stages. Earlier works suggest 10°C, 34 °C, and 45 °C as the minimum, optimum, and maximal temperatures for melon development (Baker and Reddy 2001; Bouzo and Küchen 2012). Notably, several studies indicate that specific accessions display improved tolerance to low temperatures. The Annas Yoqna’am (AY) accession, for example, displays earlier emergence, greater carbon assimilation, and enhanced biomass accumulation under suboptimal thermal conditions relative to other melon lines (Burger et al. 2010; Paris et al. 2013), indicating that adaptive mechanisms conferring early-season vigor under climatic instability are present within the species.

Grafting is widely recognized as an effective strategy for enhancing tolerance to abiotic and biotic stresses in horticultural crops (Colla et al. 2010; Schwarz et al. 2010; Yetisir and Uygur 2010; Razi et al. 2024). Stress-tolerant rootstocks can improve the performance of sensitive scions through root–shoot signaling pathways that modulate whole-plant physiology (Martínez-Ballesta et al. 2010; Ntatsi et al. 2017; Lu et al. 2022). Root responses to environmental conditions, particularly temperature, strongly influence scion physiological and developmental behavior (Dodd et al. 2000; Ntatsi et al. 2017; Aidoo et al. 2019). It is found to be mediated by long-distance transport of hormones (Sharp and Lenoble 2002) and, more recently, by the movement of macromolecules such as mRNAs and proteins across the graft junction (Kragler and Bock 2025; Zhang et al. 2025). Although grafting has demonstrated clear benefits for managing soil-borne diseases and improving tolerance to abiotic stresses in cucurbits, including melon (Aloni et al. 2010; Colla et al. 2010; Parthasarathi et al. 2021; Lu et al. 2022; Razi et al. 2024), most studies have relied on interspecific combinations and have not fully explored reciprocal influences between graft partners.

Climate-driven shifts in cropping schedules increasingly push warm-season crops such as melon into cooler early-season environments to avoid late-season heat, high vapor pressure deficits, and pathogen pressure (Eastham et al. 1999; Li et al. 2024). While such shifts may reduce exposure to extreme stress, they also expose thermophilic crops to suboptimal temperatures during emergence and early growth, often resulting in reduced vigor and delayed establishment. Melon In this context, grafting offers an immediate and flexible complement to traditional breeding, enabling rapid enhancement of plant performance under adverse conditions (Snowdon et al. 2021).

In the present study, we addressed this challenge by employing reciprocal grafting between melon accessions with contrasting responses to suboptimal temperatures. By evaluating a suite of physiological and morphological traits across three controlled temperature regimes, we sought to disentangle the individual contributions of the scion and rootstock to the overall plant response to thermal stress. This approach enables a more mechanistic understanding of graft-mediated resilience and identifies accession-specific traits that may support early-season vigor under increasingly variable climatic conditions.

## 2. Material and methods

### 2.1 Plant material and growing conditions

Two melon accession were identified from a larger collection (Gur et al., 2017), with low and high tolerance for low temperatures, PI414723 (Origin – India) and Annas Yoqna’m -AY (Origin – Israel), respectively (Burger et al. 2010; Paris et al. 2013). To examine the effect of accessions as both rootstock and scion, reciprocal grafting combinations were performed at the Hishtil nursery (Ashkelon, Israel), along with non-grafted plants, resulting in six total combinations: AY, PI414723, A/A (AY/AY), A/P (AY/PI414723), P/A (PI414723/AY), and P/P (PI414723/PI414723), representing scion/rootstock, respectively. After healing of grafted area, both grafted and non-grafted seedling were transplanted in two liter pots with local soil (chromic Haploxerert, fine-clayey, montmorillonitic, thermic), with 55% clay, 25% silt, and 20% sand; 2% organic matter; and a pH of 7.2.

Plant were grown in three growing chambers with three constant temperature regimes: 16 °C, 25 °C, and 35 °C representing the early spring sub optimal, ambient and extreme temperatures. All chambers had uniform artificial PAR lights (400-700 nm, ∼1000 µmol m^−2^ s^−1^, 12/12 hours day/night period), and constant relative humidity of 40%. In each temperature regime each combination were grown in 15 replicates. To characterize plant growth dynamics, three measurement campaigns were conducted under each temperature regime, corresponding to the following phenological stages: two–three, three–four, and four–five true leaves. At each campaign, five replicates per rootstock\scion combination were samples.

### 2.2 Physiological and phenological measurements

Starting two weeks after transplanting, plants were sampled weekly over three weeks period. The dynamic sampling included physiological and morphological parameters. The physiological sampling included leaf-level CO₂ and H₂O flux measurements, along with plant pulse amplitude modulation (PAM) measurements, that were conducted using a Licor 6800F (Licor, Lincoln, NB, USA) under light conditions. Measurements were taken for carbon assimilation rate (*An*), stomatal conductance (*g_sw_*), and photosystem II quantum yield of photochemistry (*Y(Ⅱ)*; Genty et al., 1989). For the morphological sampling, the foliage was removed close to the soil level. Afterward, the soil was removed from the root system using a gentle water stream over a sieve. The above- and below-ground biomass were then oven-dried at 72 °C for 48 hours and weighed.

At the last campaign (third round), further morphological and physiological parameters were collected. Under dark conditions, former to lights activation, dark respiration (*Rn*) and photosystem II functionality (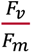; Murchie & Lawson, 2013) were measured. Additionally, the rapid A/Ci curve method (Stinziano et al. 2019) was applied, and maximum carboxylation rate (*V_cMax_*) and maximum electron transport rate (*J_Max_*), were estimated. Finally, before oven drying, root systems were carefully cleaned of soil particles and any unrelated vegetative material. The cleaned roots were then scanned and analyzed using the WinRhizo system (Regent Instruments Inc., Ottawa, ON, Canada) to quantify root length and diameter.

### 2.3 Statistical analysis

To distinguish the effects of scion and rootstock origin from the grafting procedure itself, non-grafted plants and reciprocal grafted plants were analyzed separately. The experiment was held in complete randomize design with 15 reputations per each treatment. For comparisons between the parental accessions (AY and PI414723), data normality was assessed using the Shapiro–Wilk test, and variance homogeneity was evaluated with Levene’s test. Then, differences between accessions were analyzed using one-way analysis of variance (ANOVA), followed by either Student’s t-test (non-grafted) or Tukey’s HSD test (reciprocal grafting) with Bonferroni correction for multiple comparisons (α < 0.05). These analyses were conducted using JMP® Pro 17 (SAS Institute, Cary, NC).

The reciprocal grafted population was analyzed independently of the non-grafted accessions to specifically evaluate the contributions of scion and rootstock without confounding effects from the grafting process. Where necessary, data were transformed (logarithmic, Johnson Su, or Box–Cox transformations) to improve normality and homoscedasticity prior to analysis. When transformation failed to achieve normality, the data were excluded from analysis (e.g., 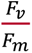 and shoot dry weight). A two-way linear model was applied in JMP® Pro 17 to assess the effects of temperature (temp) and grafting treatment (graft), along with their interaction. Specifically, the linear model was:

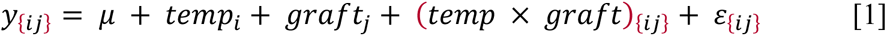

where μ is the overall mean, *temp*_*i*_is the effect of temperature level i, *graft*_*j*_ is the effect of grafting treatment j, (temp × graft){ij} is the interaction, and ε{ij} is the random error. Model fit was evaluated based on R^2^ values and p-values obtained from the “Fit Model” platform, and significant differences among least-squares means were determined using Tukey’s HSD test. In addition, comparisons between self-grafted and non-grafted plants (AY vs. A/A and PI414723 vs. P/P) were conducted to evaluate the effect of grafting on plant responses to temperature, following the same statistical approach as used for the reciprocal grafted population.

To integrate the large number of measured traits, including physiological and phenological variables from both shoot and root systems, a principal component analysis (PCA) was performed using the FactoMineR package in R. Loading factors were extracted to determine the contribution of each trait to the principal components, enabling assessment of the overall effects of temperature and grafting treatments.

## 3. Results

### 3.1 Low temperature emphasizes the differences between parental accessions

Comparisons between the non-grafted plants revealed pronounced differences under suboptimal temperature conditions (Figure 1, Table S1). Physiological traits such as carbon assimilation (*A_n_*: 7.45 vs. 3.21 µmol m⁻² s⁻¹ CO₂), stomatal conductance (g_sw_: 0.17 vs. 0.05 mol m⁻² s⁻¹ H₂O), quantum yield of photochemistry (Y(II): 0.12 vs. 0.08), and photosystem II efficiency (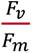: 0.54 vs. 0.38) differed significantly between the AY and PI414723 accessions, respectively. The tolerant accession (AY) also exhibited greater shoot biomass (0.53 vs. 0.43 g) and root length (616 vs. 197 cm) under low temperature. At the optimal temperature, differences between accessions were limited to root diameter (1.02 vs. 0.38 mm) and length (1134 vs. 473 cm) in AY. Under high temperature (35°C), PI414723 showed higher carbon assimilation (9.05 vs. 5.55 µmol m⁻² s⁻¹ CO₂) and lower respiration rate (*R_n_*: −2.22 vs. −4.5 µmol m⁻² s⁻¹ CO₂) than AY.

**Figure 1.**
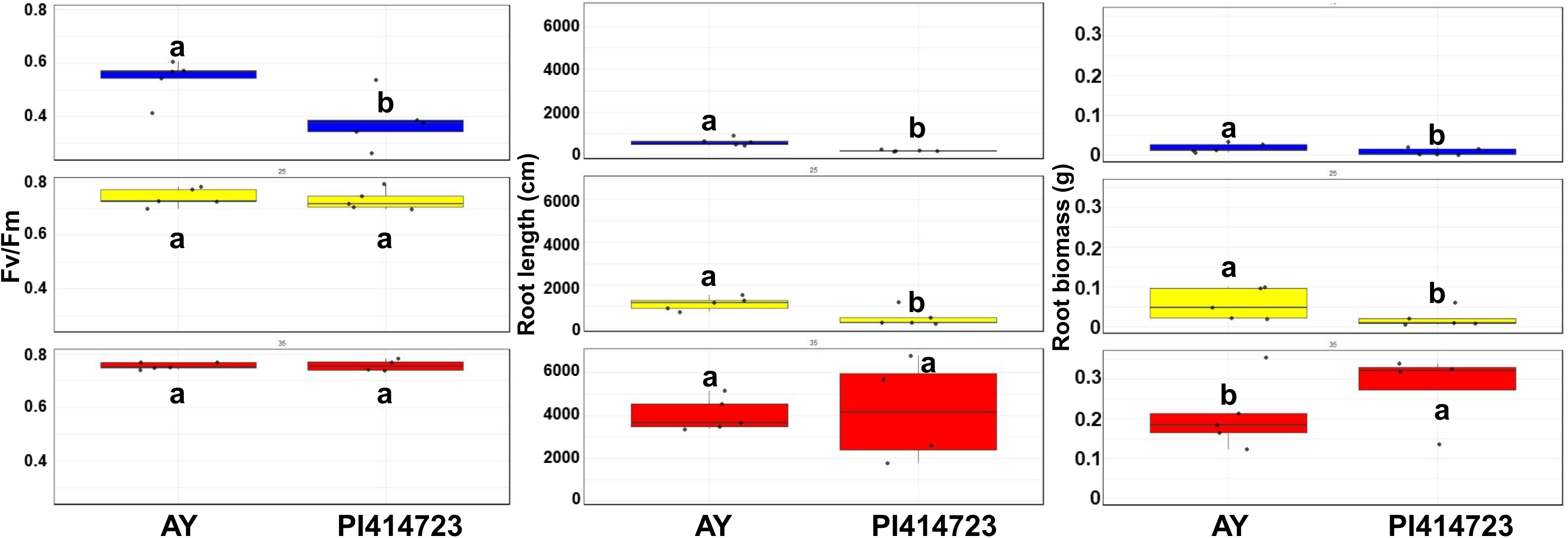
Effects of temperature regimes on PSⅡ efficiency (left column), root length (middle column), and root biomass (right column) in temperature-tolerant (AY) and temperature-susceptible (PI414723) melon accessions. Temperature treatments are color-coded: 16 °C (blue), 25 °C (yellow), and 35 °C (red). Data points represent individual plants; box plots show the mean (horizontal line) and standard deviation (whiskers). Different letters indicate statistically significant differences based on Student’s t-test (α < 0.05).

Moreover, under 16 °C the AY and PI414723 were different over the three measurement campaigns, and the AY carbon assimilation rate was approximately two times higher than the PI414723 (Figure 2a). Under 25 °C, both accessions demonstrated a similar *A_n_* rates over the three measurement campaigns (Figure 2b). unlike, under 16 °C and 25 °C, the *A_n_* values under 35 °C decline from 16 and 12 µmol m^−2^ s^−1^ in the first measurement campaign to 5.5 and 6.8 µmol m^−2^ s^−1^ for AY and PI414723, respectively (Figure 2c).

**Figure 2.**
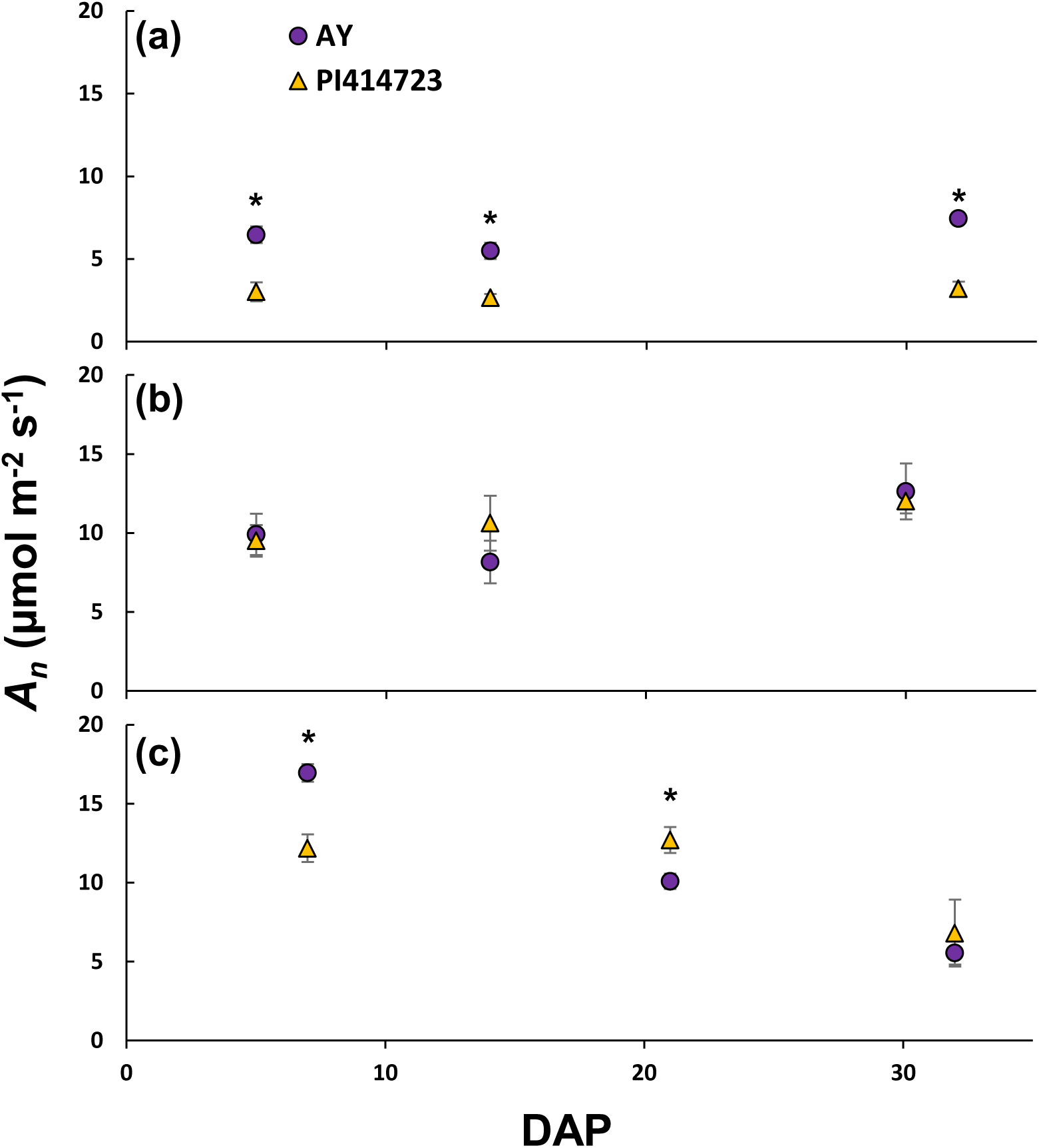
Carbon assimilation rates in the suboptimal temperature–tolerant AY and the susceptible PI414723 melon accessions under temperature regimes of 16 °C (a), 25 °C (b), and 35 °C (c). Values represent means ± SE (n = 5). Asterisks indicate statistically significant differences between accessions at the same time point based on one-way ANOVA. DAP, days after planting.

### 3.2 Phenology is the main component distinct between temperatures

Principal component analysis (PCA) of non-grafted plants confirmed temperature as the major driver of variance (Figure 3). PC1 explained 53.2% and PC2 21.1% of the total variance, with physiological and phenological parameters contributing ∼60% and ∼40% respectively to PC1 (Table 1). PC2 was dominated by physiological traits (∼94%). Temperature-specific PCA analyses further supported consistent differences between accessions across all regimes (Figure S1), with physiological traits dominating PC1 under 16°C (∼73%) and 35°C (∼78%), and to a lesser extent at 25°C (∼60%) (Tables S2).

**Figure 3.**
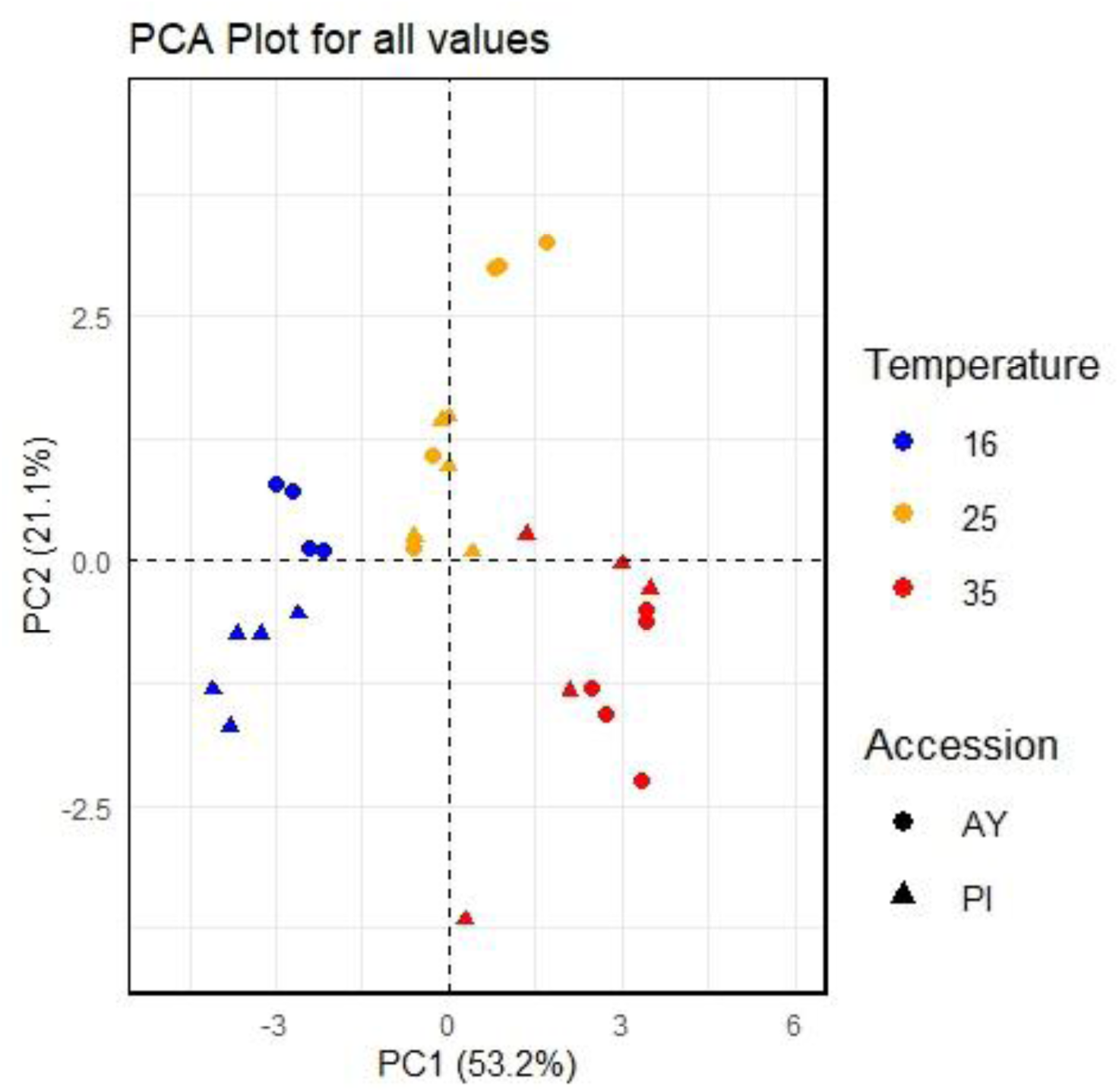
Principal Component Analysis (PCA) of the measured parameters in temperature-tolerant (AY-AY) and susceptible (PI414723-PI) melon accessions. The first and second letters represent the scion and rootstock, respectively. The plot displays the distribution of samples along the first two principal components, PC1 (53.2%) and PC2 (21.1%). Colors represent different temperature treatments (16°C, 25°C, and 35°C), while shapes correspond to the melon accessions. Dashed lines indicate the origin of the principal components.

**Table 1.**
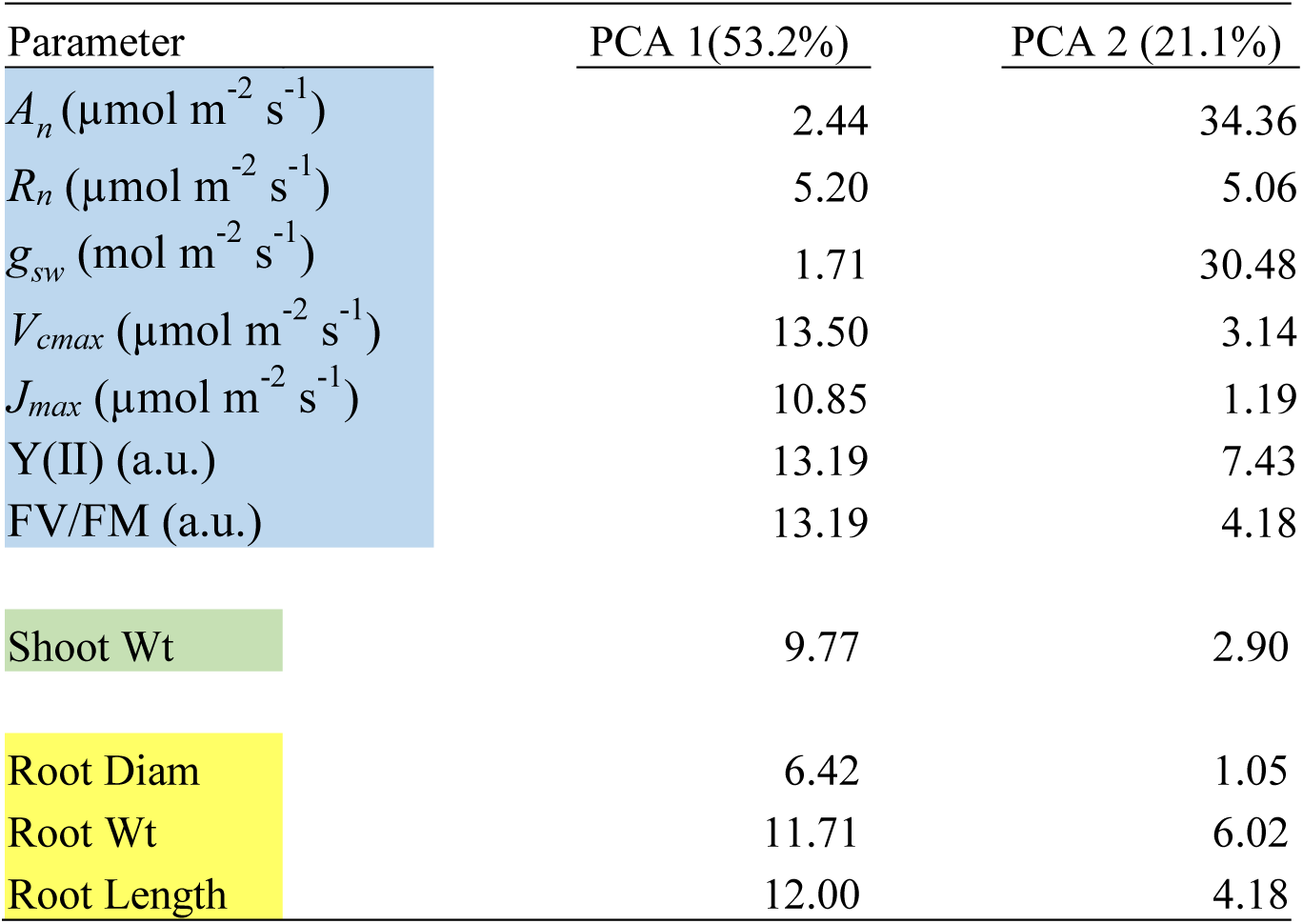
Loadings of Principal Component 1 (PC1) and Principal Component 2 (PC2) for all measured parameters between Parent accessions. Parameters are categorized into leaf-level physiological traits (Blue), shoot biomass (Green), and root traits (Yellow).

Grafted plants also showed strong responses across temperature regimes (Table S3). PCA conducted on all grafted combinations revealed PC1 and PC2 explained 49.9% and 25.7% of the variance, respectively (Figure 4). PC1 primarily separated the temperature regimes, while PC2 differentiated 25°C from 16°C and 35°C. Phenological traits accounted for 54.25% of the variability in PC1, whereas physiological traits dominated PC2 (94.75%) (Table 2).

**Figure 4.**
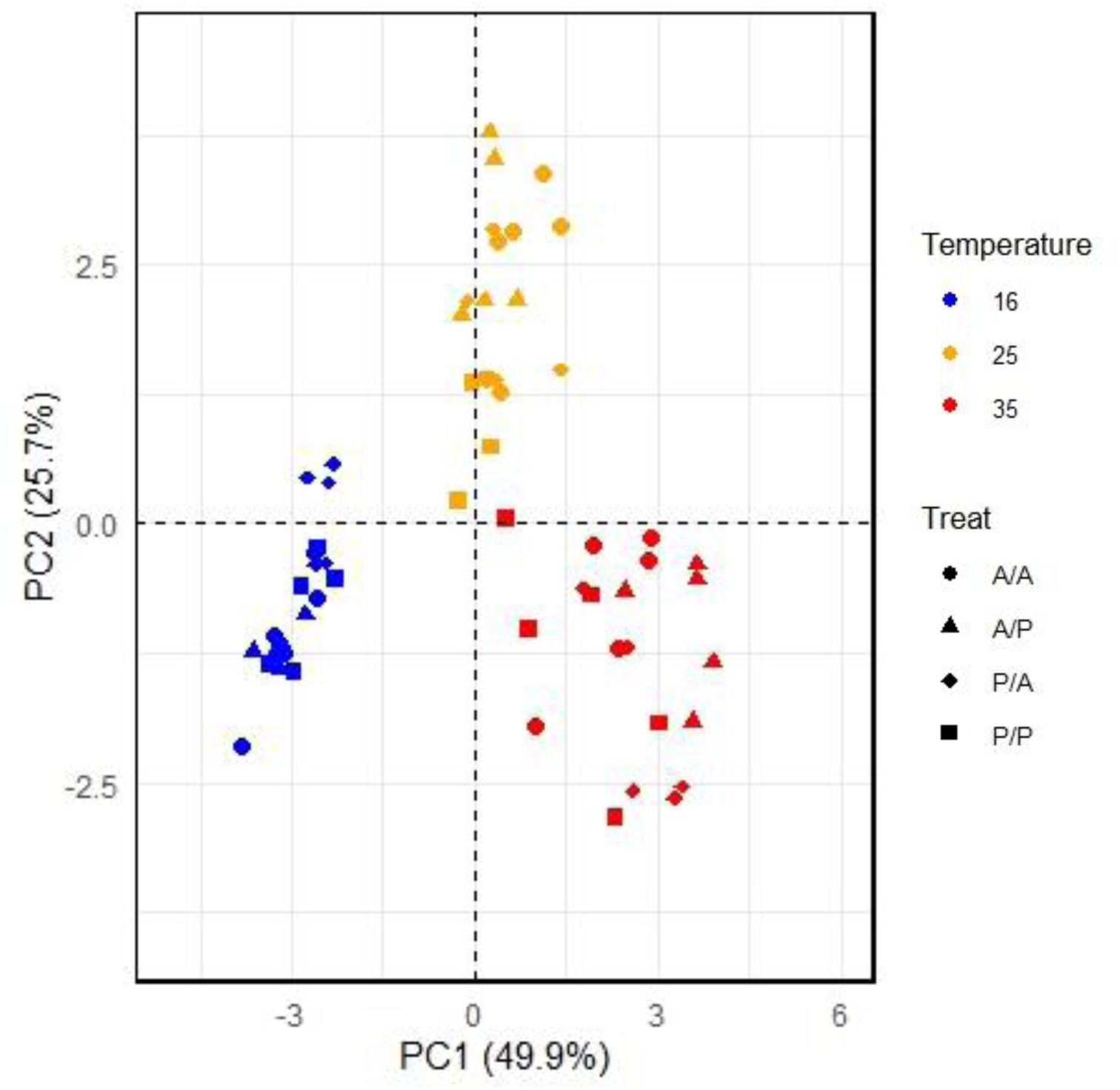
Principal Component Analysis (PCA) of the measured parameters in grafted plants from temperature-tolerant (AY-A) and susceptible (PI414723-P) melon accessions. The first and second letters represent the scion and rootstock, respectively. The plot displays the distribution of samples along the first two principal components, PC1 (49.9%) and PC2 (25.7%). Colors represent different temperature treatments (16°C, 25°C, and 35°C), while shapes correspond to the reciprocal grafting treatments (A/A, A/P, P/A, and P/P). Dashed lines indicate the origin of the principal components.

**Table 2.**
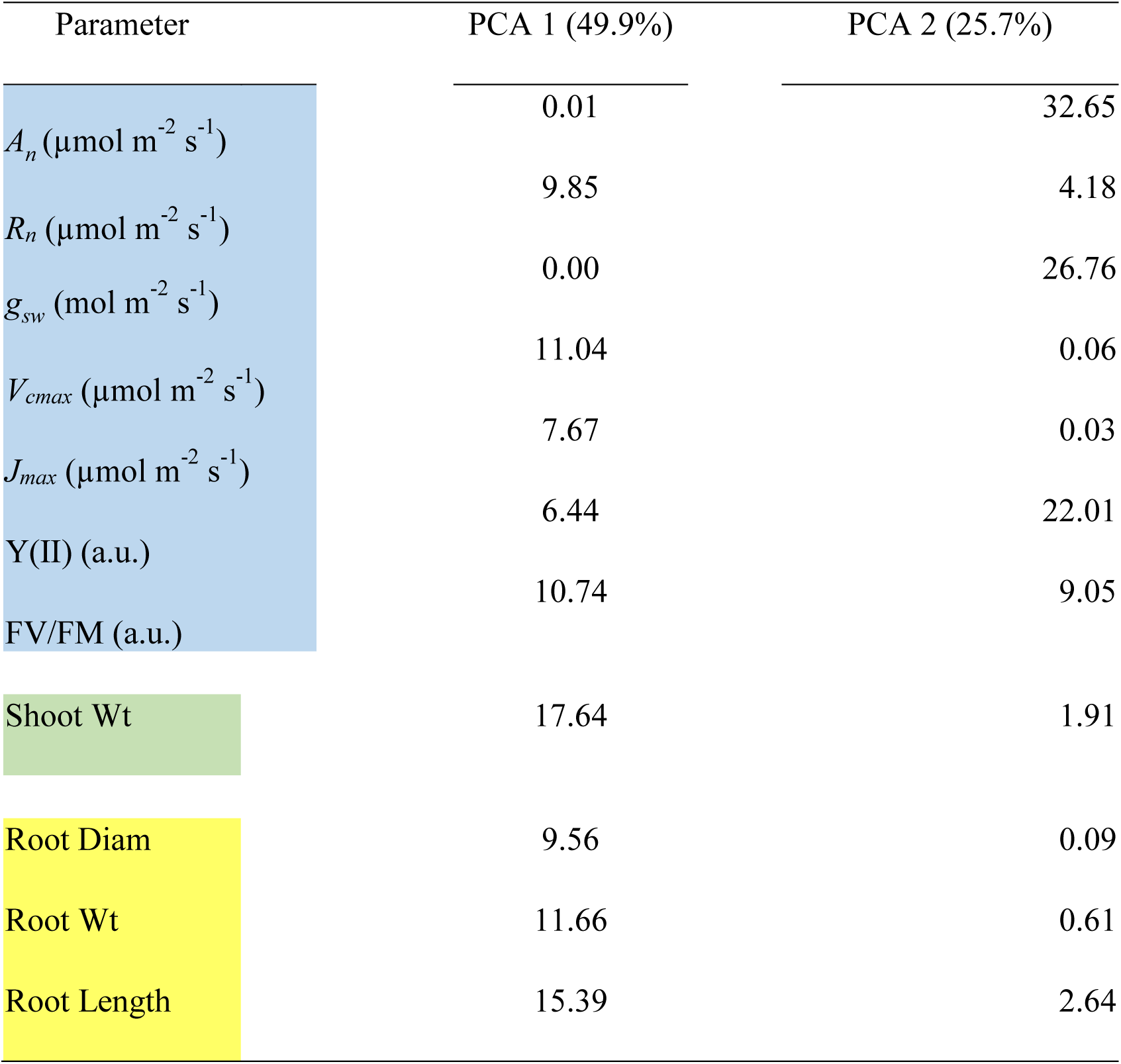
Loadings of Principal Component 1 (PC1) and Principal Component 2 (PC2) for all measured parameters across all temperature regimes in the reciprocal grafted plants. Parameters are categorized into leaf-level physiological traits (Blue), shoot biomass (Green), and root traits (Yellow).

### 3.3 Scion is the dominant part in low temperature response

Given the strong effect of temperature, separate PCAs were conducted for each regime (Figure 5). In all cases, PC1 and PC2 explained over 50% of the variance (Tables 3). PC1 was largely driven by physiological traits (>80%), and PC2 by phenological traits, particularly at 25 °C and 35 °C (54% and 69%, respectively), but less so at 16 °C (36%). At 16 °C, PC1 clearly separated treatments based on scion identity (Figure 5a), whereas no clear scion/rootstock-based separation was observed at 25 °C and 35 °C (Figures 5b, 5c). However, at 25 °C, distinct clustering was seen: A/A and A/P separated along PC2, while P/P and P/A diverged along PC1. Fitted models confirmed that all measured traits, except *R_n_* and *J_max_*, were significantly influenced by both grafting and temperature (Table 4). All models explained more than 59% of the variance of the measured traits.

**Figure 5.**
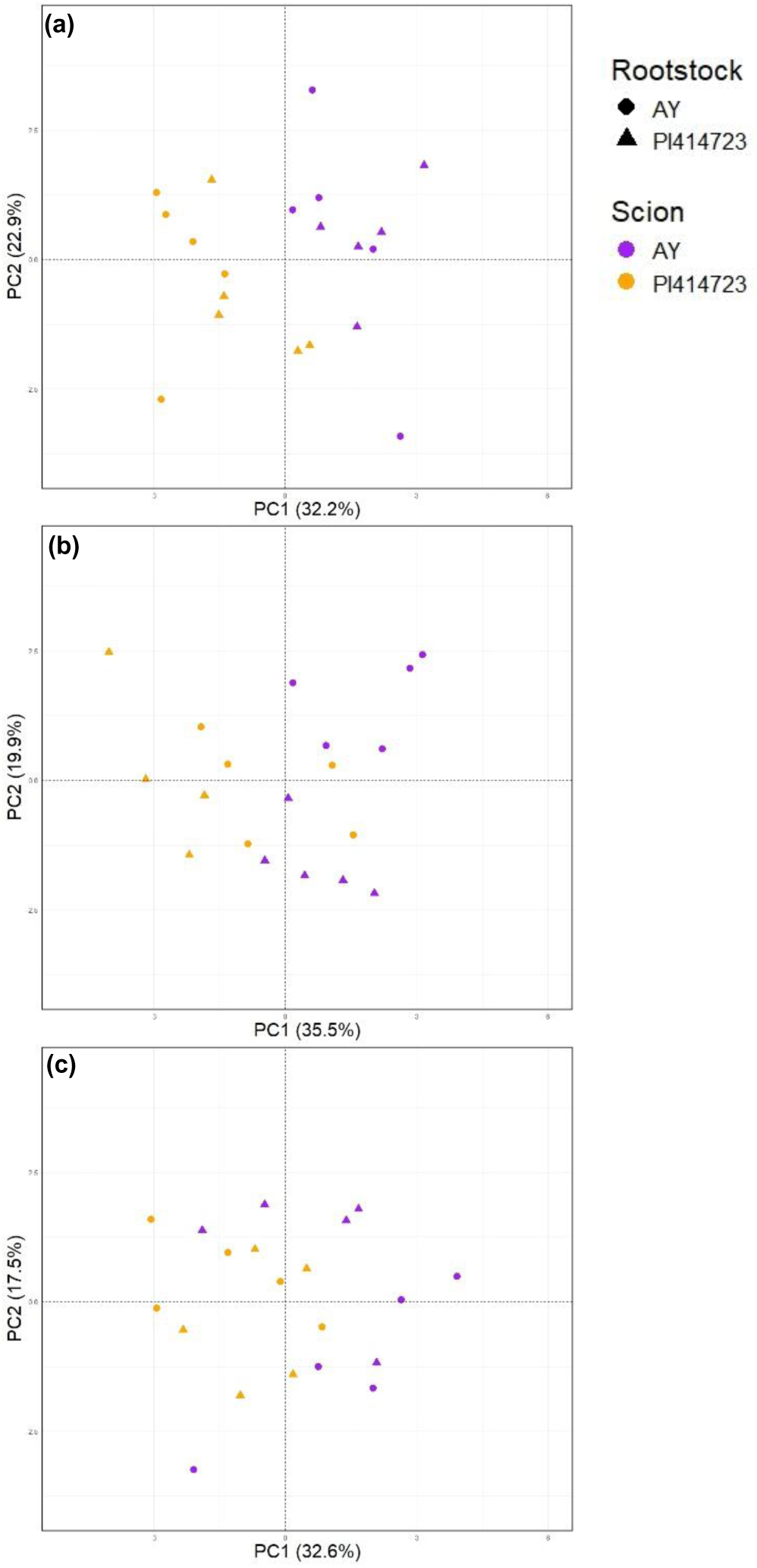
Principal Component Analysis (PCA) of the measured parameters in grafted plants from temperature-tolerant (AY) and susceptible (PI414723) melon accessions under 16°C (a), 25°C (b), and 35°C (c). The plot illustrates the distribution of samples along the first two principal components, PC1 and PC2. Shapes represent the reciprocal grafting rootstock, while colors indicate the scion. Dashed lines mark the origin of the principal components. Loading parameters for PC1 and PC2 at 16°C, 25°C, and 35°C are provided in Table 3.

**Table 3.**
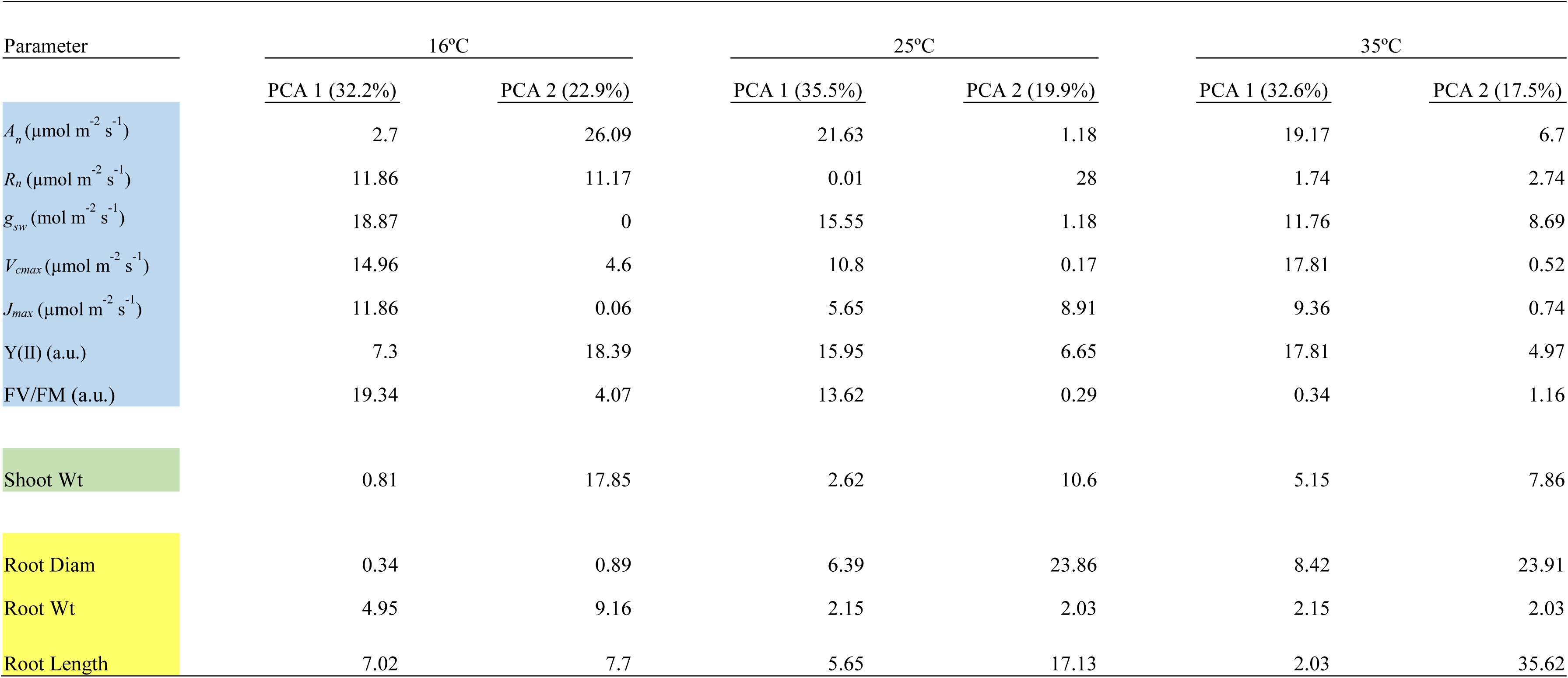
Loadings of Principal Component 1 (PC1) and Principal Component 2 (PC2) for all measured parameters in the grafted plants under 16°C, 25°C, and 35°C. Parameters are categorized into leaf-level physiological traits (Blue), shoot biomass (Green), and root traits (Yellow).

**Table 4.**
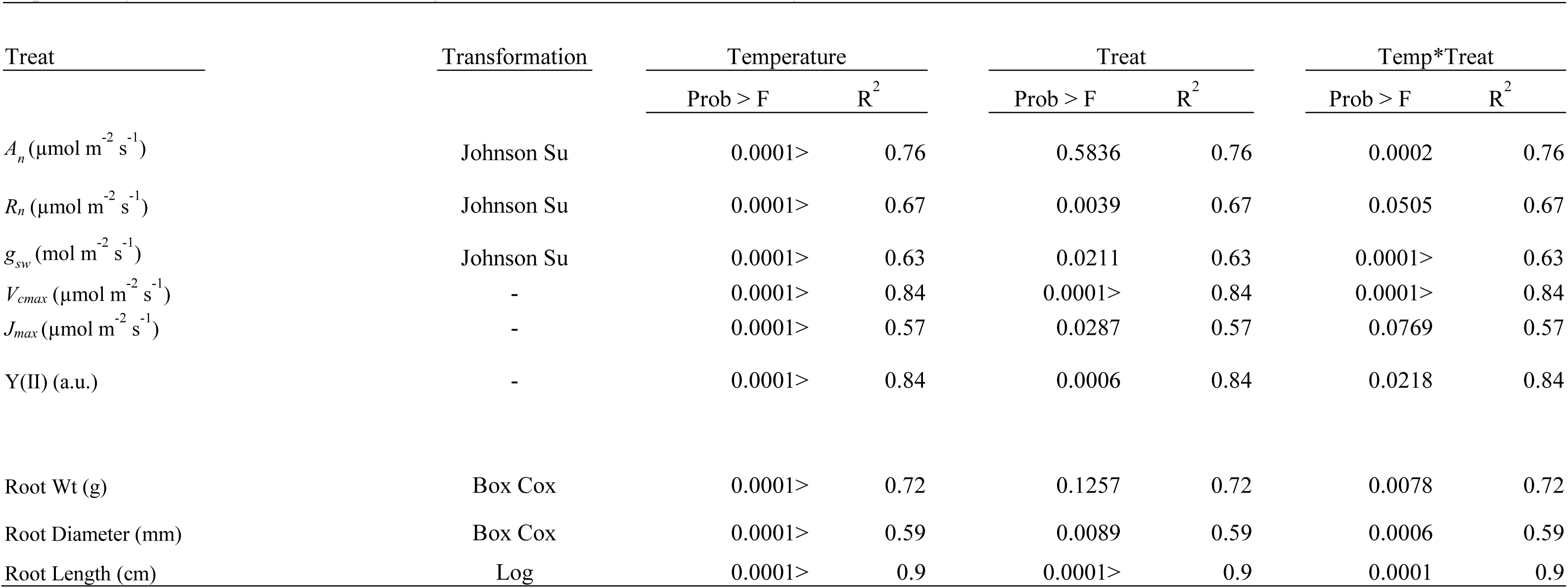
The effects of temperature, grafting, and their interaction on the measured traits. Data were transformed where necessary to meet normality assumptions. A probability value below 0.05 indicates a significant difference based on the Tukey HSD test.

### 3.4 Sustained high temperatures eventually impair physiological function

Carbon assimilation followed distinct temporal patterns across temperature treatments (Figure 6). At 16 °C, assimilation rates remained low but exhibited a slight upward trend over time. At 25 °C, rates progressively increased throughout the measurement period. In contrast, at 35 °C, all plant types experienced a decline in assimilation over time. The A/A combination showed the steepest decline, decreasing from 17.8 to 5.7 µmol m⁻² s⁻¹ CO₂, while P/P plants showed a milder reduction from 14.2 to 5.8 µmol m⁻² s⁻¹ CO₂. Statistically significant differences among the grafted treatments were detected at several measurement points; however, these effects did not exhibit a consistent pattern across time or traits (Table S3). Other physiological parameters, such as stomatal conductance (gsw) and quantum yield of photochemistry (Y(II)), displayed comparable trends (Table S4).

**Figure 6.**
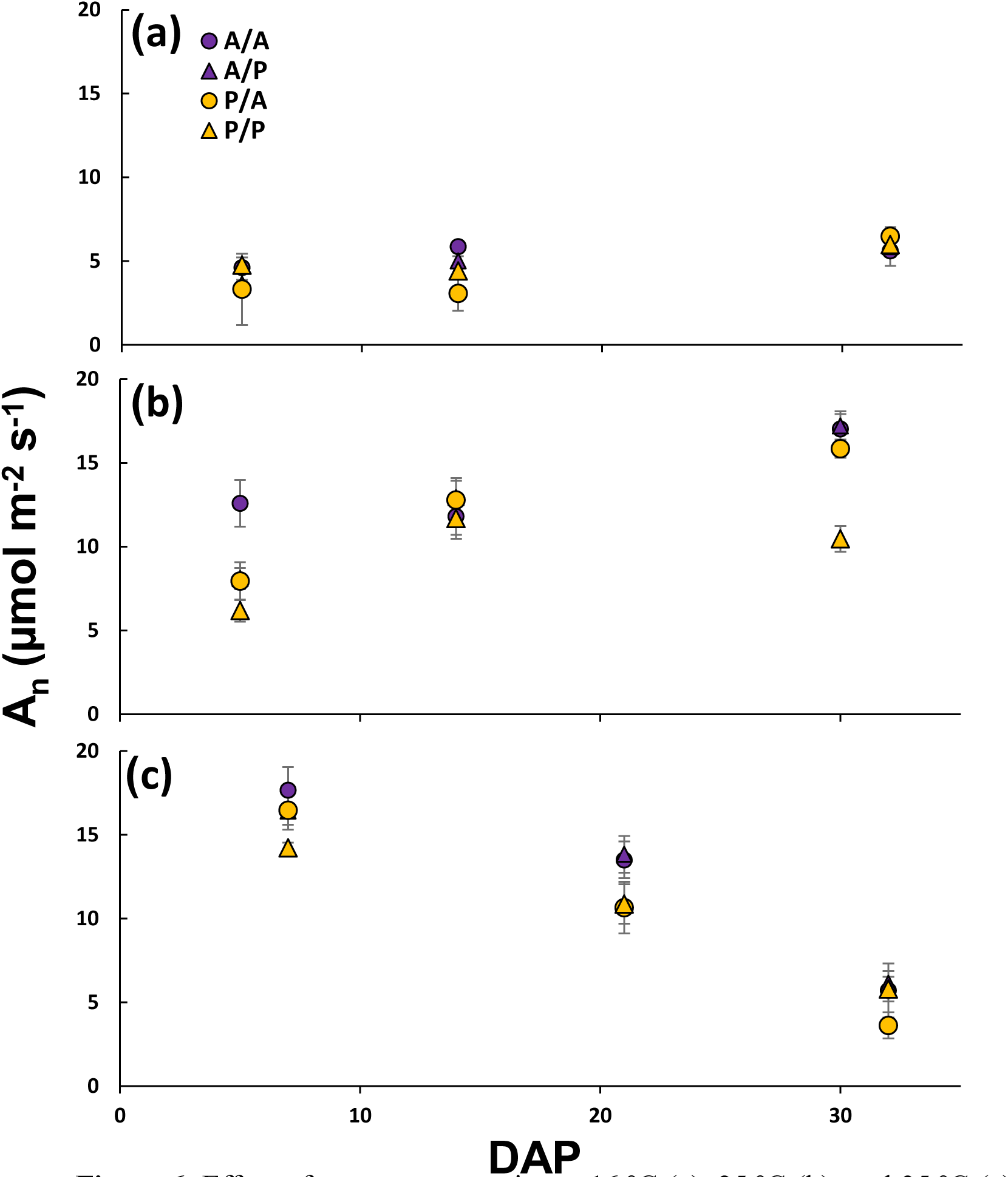
Effect of temperature regimes: 16 °C (a), 25 °C (b), and 35 °C (c), on the carbon assimilation rate of A/A (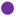), A/P (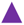), P/A (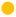), and P/A (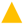). Each point represents the mean of five plants, and error bars indicate the standard error. DAP, days after planting. Statistical analysis of each time point are attached in table S3.

### 3.5 Grafting did not alter the physiological functioning of the plant

Comparison between non-grafted and self-grafted plants (AY vs. A/A, PI414723 vs. P/P) revealed minimal physiological differences (Tables S5–S6). Apart from a *J_max_* difference in AY-A/A and a grafting-temperature interaction affecting *A_n_* in PI414723-P/P, no other significant grafting effects were observed.

## 4. Discussion

In this study, we aim to investigate the effects of low temperature-susceptible and -tolerant melon rootstocks and scions under non-optimal temperature conditions, with the goal of enhancing crop adaptation to early-season sowing and unexpected cold events. The reciprocal grafting of the plants reveal that the main contribution to suboptimal temperatures tolerance is the scion. The contribution of the tolerant scion was pronounced not only in the scion measured parameters but also in the rootstock traits as root length and weight. Both row data and PCA analysis indicate that the main source for the difference between treatments is the biochemical activity. It is well established that plant biochemical processes are strongly temperature dependent (Bernacchi et al. 2002, 2003). In this study, differences in the maximum carboxylation capacity of Rubisco (*Vc_max_*) and the maximum electron transport rate supporting RuBP regeneration (*Jₘₐₓ*) help explain the superior carbon assimilation efficiency observed under cooler conditions. Higher *Vcₘₐₓ* and *Jₘₐₓ* values support more effective Calvin-cycle activity and reduce the need for excess energy to be dissipated through alternative metabolic pathways (Zait et al. 2024). Furthermore, the elevated mitochondrial respiration rates (*Rn*) observed in P/A and P/P plants suggest that, although net CO₂ assimilation (*Aₙ*) appears similar among the graft combinations, a larger fraction of the assimilated carbon is consumed by respiration in these grafted plants. Consequently, the actual carbon available for growth and biomass accumulation is reduced (McAusland et al. 2023).

The sustained physiological activity observed in the scion at low temperatures found to support root development, potentially through the translocation of regulatory compounds. Previous studies have emphasized the movement of molecules such as hormones, mRNAs, and proteins from root to shoot to maintain shoot function during stress (Aidoo et al. 2019; Lu et al. 2022; Li et al. 2023; Kragler and Bock 2025; Zhang et al. 2025). Our findings suggest that signaling from shoot to root may also contribute to the induction of stress tolerance belowground. Hormones as ABA are known to promote both stomatal closure and inhibit root elongation (Honour et al. 1995; Rowe et al. 2016). In interaction with other hormones in the foliage, it can control the plant belowground response (Sharp and Lenoble 2002; Tiwari et al. 2023) but further research is required.

Many studies have shown that tolerant rootstock can contribute to the susceptible scion’s response to low temperatures (Venema et al. 2008; Lu et al. 2022; Razi et al. 2024). These studies address tolerant rootstock’s impact on susceptible scions, but not their reciprocal effect. Moreover, these studies focuses on grafting a susceptible crop as a scion onto a dominant rootstock, which may obscure the mutual effects of the grafting interaction. This study provides evidence that grafting a temperature-tolerant scion onto a susceptible rootstock can enhance rootstock performance under suboptimal temperatures in melon.

The dominant influence of the scion under low-temperature conditions was not evident at higher temperatures in grafted plants, even though the non-grafted parental accessions showed significant differences under those same conditions. At moderate temperatures (25 °C), reciprocal grafts (P/A and A/P) were distinguishable from their respective self-grafted controls (P/P and A/A), indicating a potential interaction between graft identity and thermal response. These findings highlight that the contribution of graft components may vary across environmental conditions (Lu et al. 2022).

Given that melon was domesticated in warm climates (Endl et al. 2018), 35 °C may not constitute an extreme condition for most accessions, although, prolonged exposure to high temperatures was found to suppress plant responses over time. The broad genetic diversity within the species, including its wild relatives, increases the likelihood of identifying genotypes with enhanced tolerance to suboptimal temperatures (Sebastian et al. 2010; Paris et al. 2013). Previous work has already reported accessions with relative tolerance for low temperatures (Burger et al. 2006; Paris et al. 2013). In the context of climate change and shifting cultivation calendars, our study aimed to clarify whether tolerance arises from the scion or rootstock and whether it can be conferred through grafting. In future research, we will extend this framework to explore the response of these accessions to supra-optimal temperatures.

The PCA analysis of the full dataset revealed that temperature plays a major role in both physiological and morphological responses. Biomass accumulation in the shoot, and in most root samples, was highest under the warmest temperature conditions. However, the physiological responses were more pronounced. Plants grown at 35 °C initially exhibited the highest carbon assimilation rates initially, but these declined sharply at subsequent measurement campaigns. In contrast, plants at the moderate temperature (25 °C) showed a gradual increase in carbon assimilation over time. Although elevated temperatures are known to enhance photosynthetic activity and carbon assimilation in plants (Bernacchi et al. 2003), prolonged exposure to high temperatures can have detrimental effects. While melons are generally well-adapted to high temperatures, extended exposure, such as applied in this study, can eventually reduce physiological performance and accelerate senescence (De la Haba et al. 2014). The accumulation of reactive oxygen species (ROS) can cause cellular damage, ultimately leading to cell death and early senescence (Zhao et al. 2011). These results indicate that while extreme heat maximizes short-term growth, moderate temperatures support more sustained physiological performance.

Comparisons between the non-grafted accessions revealed significant physiological and morphological differences, under all temperature. These results are in agreement with (Burger et al. 2010; Paris et al. 2013), which identified two melon accessions with distinct physiological and phenological responses to temperature variations. However, the current study revealed that the main source of difference between the accessions was their physiological more than their phenological response under the different temperatures. Further, we observed that under the low temperature, in addition to the physiological differences, morphological changes were found. Both shoot biomass, and root traits such as length, and biomass accumulation were significantly higher in the AY tolerant accession under these low temperatures. However, under optimal and high temperatures almost no differences were observed between the two accessions. These results highlight that melon plants, due to their origin, are generally adapted to high temperatures. Moreover, adaptation to low temperatures is uncommon despite the large variety within this plant species (Gur et al., 2017).

Grafting was suggested as an effective method to increase the plant vigor, biotic and abiotic stress tolerance in addition to other positive effect on the scion plant behavior (Venema et al. 2008; Yetisir and Uygur 2010; Simpson et al. 2015; Dafna et al. 2021; Parthasarathi et al. 2021). Due to that, we decided to apply a reciprocal grafting to investigate the cross talk between the scion and rootstock under unfavorable temperature response in addition to self-grafting as a control. Comparison of self-grafted plants with their non-grafted counterparts (PI414723-P/P vs. PI414723 and AY-A/A vs. AY) revealed minimal effects of grafting on plant physiology. Apart from a significant grafting effect on *J_max_* in AY-A/A and an interaction between grafting and temperature on carbon assimilation in PI414723-P/P, no other major effects were detected. These results suggest that while grafting may influence specific physiological traits, it does not significantly alter the overall response to temperature.

In the current study, each temperature regime was applied as a constant condition to isolate its specific effects. However, it is important to note that separating day and night temperatures, even if the mean remains the same, can reduce plant stress levels and potentially lead to different physiological outcomes. This highlights the importance of considering temperature fluctuations when interpreting plant responses under controlled conditions (Sunoj et al. 2016).

## 5. Conclusions

This study demonstrates that reciprocal grafting between suboptimal temperature-tolerant and susceptible melon accessions, highlight the often-overlooked contribution of scion traits to abiotic stress tolerance in grafted plants. Although rootstocks are commonly viewed as the primary drivers of abiotic stress tolerance, our results show that scion-derived biochemical properties, particularly those governing Calvin cycle activity, strongly influence plant performance under low temperatures. A tolerant scion enhanced root system elongation even when grafted onto a susceptible rootstock, whereas a susceptible scion constrained the performance of a tolerant root system. These findings highlight the bidirectional nature of root–shoot interactions and suggest that optimizing scion traits is as important as rootstock selection for improving crop adaptation to cooler environments. Incorporating scion-driven physiological mechanisms into breeding strategies may support the development of cultivars better suited to diverse climatic regions and future climate-change scenarios.

## Acknowledgement

The authors wish to thank to Ofir Tal, and Elad fein for help at various stages of the project.

## Conflict of interest

The authors declare no conflicts of interest.

## Data availability

Data will be made available on reasonable request.

## Notes

### Competing Interest Statement

The authors have declared no competing interest.

