## supplemental data for "Physiological dominance of the scion in shaping root architecture under suboptimal temperature"

**Table S1**. The impact of growth temperatures on physiological and phenological traits in two melon accessions: temperature-tolerant (AY) and susceptible (PI414723). Values represent the mean ± standard deviation of five replicates. Different letters denote statistically significant differences between accessions under the same temperature condition, as determined by Student's t-test ($\alpha < 0.05$).

| | 16ºC | | 25ºC | | 35ºC | |
|---|---|---|---|---|---|---|
| Trait | AY | PI414723 | AY | PI414723 | AY | PI414723 |
| $A_n$(µmol $m^{-2}$ $s^{-1}$) | 7.45±0.38a | 3.21±1.01b | 12.64±4.42a | 12.03±1.96a | 5.55±1.87b | 9.05±1.94a |
| $R_n$ (µmol $m^{-2}$ $s^{-1}$) | -1.91±0.85a | -1.7±1.1a | -1.96±0.43a | -1.92±0.61a | -4.5±0.83b | -2.22±0.14a |
| $g_{sw}$ (mol $m^{-2}$ $s^{-1}$) | 0.17±0.04a | 0.05±0.08b | 0.26±0.09a | 0.16±0.07a | 0.13±0.06a | 0.18±0.06a |
| VcMax (µmol $m^{-2}$ $s^{-1}$) | 17.52±6.6a | 29.43±24.4a | 61.54±14.17a | 82.06±11.98a | 110.21±12.21a | 106.93±12.68a |
| Jmax (µmol $m^{-2}$ $s^{-1}$) | 33.07±22.49a | 45.62±19.13a | 111.98±51.40a | 96.6±21.43a | 144.68±20.68a | 129.1±45.22a |
| Y(II) (a.u.) | 0.12±0.01a | 0.08±0.02b | 0.32±0.1a | 0.25±0.03a | 0.32±0.05a | 0.26±0.07a |
| Fv/Fm (a.u.) | 0.54±0.07a | 0.38±0.10b | 0.74±0.03a | 0.73±0.04a | 0.76±0.01a | 0.76±0.02a |
| Shoot Wt (g) | 0.53±0.05a | 0.43±0.04b | 0.93±0.14a | 2.46±1.57a | 2.55±0.55a | 3.34±0.36a |
| Root Wt (g) | 0.02±0.01a | 0.01±0.01a | 0.06±0.04a | 0.02±0.02a | 0.21±0.09a | 0.28±0.1a |
| Root Diameter (mm) | 0.43±0.05a | 0.47±0.06a | 1.02±0.53a | 0.38±0.08b | 1.02±0.46a | 0.81±0.12a |
| Root Length (cm) | 616.33±181.43a | 196.63±37.69b | 1134.64±316.77a | 473.77±425.6b | 4030.82±774.68a | 4205.13±2401.91a |

Under Review

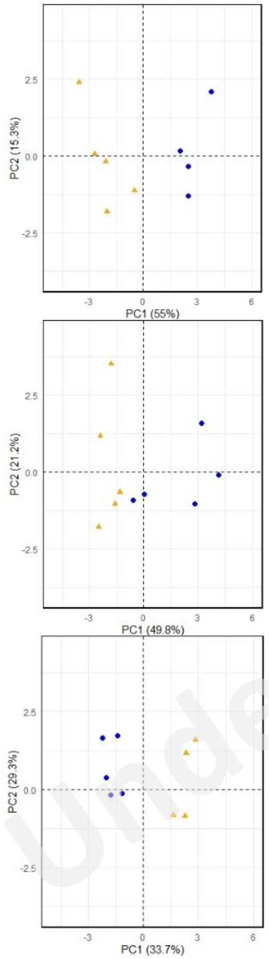


Figure S1. Principal Component Analysis (PCA) of the measured parameters in temperature-tolerant (AY - ●) and susceptible (PI414723 - ▲) melon accessions under 16ºC (a), 25ºC (b), and 35ºC (c). The plot illustrates the distribution of samples along the first two principal components, PC1 and PC2. Dashed lines mark the origin of the principal components. Loading parameters for PC1 and PC2 at 16ºC, 25ºC, and 35ºC are provided in Tables X, XX, and XXX, respectively.

Under Review

**Table S2.** Loadings of Principal Component 1 (PC1) and Principal Component 2 (PC2) for all measured parameters for AY (tolerant) and PI414723 (susceptible) under 16ºC, 25ºC, and 35ºC. Parameters are categorized into leaf-level physiological traits (Blue), shoot biomass (Green), and root traits (Yellow).

| Parameter | 16ºC | | 25ºC | | 35ºC | |
|---|---|---|---|---|---|---|
| | PCA 1 (55%) | PCA 2 (15.3%) | PCA 1 (49.8%) | PCA 2 (21.2%) | PCA 1 (33.7%) | PCA 2 (29.3%) |
| $A_n$ (µmol m$^{-2}$ s$^{-1}$) | 14.64 | 1.86 | 11.88 | 0.92 | 2.37 | 26.77 |
| $R_n$ (µmol m$^{-2}$ s$^{-1}$) | 5.5 | 11.9 | 0.8 | 36.9 | 0.24 | 27.39 |
| $g_{sw}$ (µmol m$^{-2}$ s$^{-1}$) | 10.3 | 2.28 | 10.97 | 0.69 | 12.36 | 13.03 |
| *VcMax* (µmol m$^{-2}$ s$^{-1}$) | 7.82 | 10 | 10.97 | 2.76 | 20.28 | 0.29 |
| $J_{max}$ (µmol m$^{-2}$ s$^{-1}$) | 6.09 | 12.4 | 4.06 | 6.22 | 2.54 | 4.95 |
| Y(II) | 14.32 | 0.19 | 16.96 | 3.21 | 12.75 | 0.93 |
| FV/FM | 14.64 | 0.08 | 3.88 | 10.64 | 27.8 | 0.23 |
| Shoot Wt (g) | 6.93 | 19.21 | 4.24 | 15.22 | 16.08 | 14.36 |
| Root Diam (mm) | 3.58 | 15.62 | 13.81 | 0.02 | 0 | 2.84 |
| Root Wt (g) | 5.89 | 9.11 | 13.15 | 11.06 | 4.13 | 3.71 |
| Root Length (cm) | 10.3 | 17.37 | 9.27 | 12.37 | 1.46 | 5.5 |

Under Review

**Table S3.** Effect of grafting (A = AY, P = PI414723) and temperature regime on carbon assimilation rates across three measurement campaigns corresponding to the two–three, three–four, and four–five true-leaf stages. Values represent means ± SE (n = 5). Different letters indicate statistically significant differences according to Tukey's HSD test (α < 0.05).

| Temperature | 16 | | | 25 | | | 35 | | |
|---|---|---|---|---|---|---|---|---|---|
| Round | 1st | 2nd | 3rd | 1st | 2nd | 3rd | 1st | 2nd | 3rd |
| Scion/Rootstock | | | | | | | | | |
| A/A | 4.60±0.19a | 5.87±0.36a | 5.61±0.89a | 12.6±1.40a | 11.81±1.10a | 17.03±0.90a | 17.67±1.38a | 13.5±1.10a | 5.71±1.61a |
| A/P | 3.63±0.25a | 5.01±0.28a | 5.85±0.61a | 8.1±0.63b | 12.94±1.17a | 17.2±0.88a | 16.41±1.11a | 13.84±1.09a | 6.14±0.73a |
| P/A | 3.31±1.03a | 3.07±0.56b | 6.47±0.83a | 7.95±1.14b | 12.79±1.15a | 15.86±0.54a | 16.47±0.86a | 10.65±1.54a | 3.63±0.77a |
| P/P | 4.76±0.47a | 4.41±0.30ab | 5.99±0.56a | 6.19±0.67b | 11.68±1.21a | 10.47±0.77a | 14.23±0.31a | 10.88±1.17a | 5.80±0.73a |

Under Review

Table S4. The effect of grafting (A- AY, P- PI414723) and temperature regime on the value of the different measured parameters used in the PCA analysis. Values represent the mean value±SD.

| | 16 | | | | 25 | | | | 35 | | | |
|---|---|---|---|---|---|---|---|---|---|---|---|---|
| Trait | A/A | A/P | P/A | P/P | A/A | A/P | P/A | P/P | A/A | A/P | P/A | P/P |
| Rn (µmol $m^{-2}$ $s^{-1}$) | -1.63±0.39 | -1.18±0.71 | -2.43±0.77 | -2.33±0.47 | -2.46±0.32 | -1.89±0.2 | -2.3±0.49 | -2.15±0.67 | -4.64±0.41 | -5.45±0.52 | -6.94±1.63 | -6.55±6.13 |
| $g_{sw}$ (mol $m^{-2}$ $s^{-1}$) | 0.10±0.03 | 0.07±0.07 | 0.28±0.07 | 0.11±0.05 | 0.41±0.14 | 0.29±0.1 | 0.26±0.08 | 0.19±0.05 | 0.11±0.07 | 0.15±0.04 | 0.12±0.07 | 0.13±0.05 |
| VcMax (µmol $m^{-2}$ $s^{-1}$) | 66.91±22.98 | 90.39±17.4 | 29.61±13.99 | 19.42±3.41 | 64.74±7.2 | 88.8±10.88 | 97.11±27.49 | 93.58±13.79 | 123.86±18.47 | 111.39±22.48 | 97.5±7.75 | 95.85±16.24 |
| Jmax (µmol $m^{-2}$ $s^{-1}$) | 88.64±36.42 | 98.48±32.37 | 47.09±22.2 | 73.95±45.95 | 121.35±22.43 | 106.58±24.14 | 104.85±57.53 | 135.77±64.76 | 179.96±39.73 | 147.86±18.48 | 132.04±48.7 | 93.17±16.5 |
| Y(II) (a.u.) | 0.09±0.03 | 0.11±0.02 | 0.12±0.02 | 0.11±0.02 | 0.34±0.02 | 0.39±0.04 | 0.32±0.03 | 0.27±0.05 | 0.26±0.08 | 0.3±0.06 | 0.21±0.04 | 0.22±0.06 |
| Fv/Fm (a.u.) | 0.52±0.13 | 0.45±0.06 | 0.65±0.06 | 0.55±0.11 | 0.78±0.02 | 0.8±0.01 | 0.74±0.02 | 0.71±0.01 | 0.76±0.01 | 0.77±0.01 | 0.75±0.01 | 0.77±0.01 |
| Shoot Wt (g) | 0.45±0.13 | 0.37±0.05 | 0.40±0.06 | 0.40±0.05 | 1.47±0.24 | 1.33±0.67 | 1.13±0.32 | 1.76±0.44 | 3.36±0.45 | 3.82±0.30 | 4.18±1.21 | 3.84±0.46 |
| Root Wt (g) | 0.02±0.01 | 0.02±0.01 | 0.01±0 | 0.01±0.01 | 0.12±0.05 | 0.06±0.05 | 0.24±0.28 | 0.03±0.02 | 0.14±0.09 | 0.39±0.15 | 0.34±0.11 | 0.25±0.08 |
| Root Diameter (mm) | 0.48±0.07 | 0.48±0.09 | 0.49±0.09 | 0.44±0.12 | 0.88±0.21 | 0.39±0.03 | 0.45±0.02 | 0.48±0.04 | 1.0±0.49 | 0.9±0.27 | 0.75±0.23 | 0.68±0.26 |
| Root Length (cm) | 644.78±174.34 | 428.05±178.41 | 255.72±110.57 | 206.64±108.64 | 1973.69±324.32 | 1112.89±365.8 | 1598±318.6 | 1031.53±511.52 | 4202.83±1966.23 | 7233.13±3297.06 | 7060.14±1663.58 | 4642.41±2236.91 |

Under Review

Table S5. The effects of temperature, grafting, and their interaction on the measured traits in AY accession in comparison to self grafted AY. Data were transformed where necessary to meet normality assumptions. A probability value below 0.05 indicates a significant difference based on the Student's-T HSD test.

Under Review

| Treat | Transformation | Temperature | | Treat | | Temp*Treat | |
|---|---|---|---|---|---|---|---|
| | | Prob > F | $R^2$ | Prob > F | $R^2$ | Prob > F | $R^2$ |
| $A_n$ (µmol $m^{-2}$ $s^{-1}$) | Johnson Su | 0001.> | 0.62 | 0.94 | 0.62 | 0.19 | 0.62 |
| Rn (µmol $m^{-2}$ $s^{-1}$) | Johnson Su | 0001.> | 0.71 | 0.62 | 0.71 | 0.23 | 0.71 |
| $g_{sw}$ (mol $m^{-2}$ $s^{-1}$) | Johnson Su | 0001.> | 0.67 | 0.71 | 0.67 | 0.03 | 0.67 |
| VcMax (µmol $m^{-2}$ $s^{-1}$) | - | 0001.> | 0.87 | 0.0003 | 0.87 | 0.005 | 0.87 |
| Jmax (µmol $m^{-2}$ $s^{-1}$) | - | 0001.> | 0.69 | 0.01 | 0.69 | 0.33 | 0.69 |
| Y(II) (a.u.) | - | 0001.> | 0.79 | 0.34 | 0.79 | 0.26 | 0.79 |
| Root Wt (g) | Box Cox | 0.0003 | 0.54 | 0.51 | 0.54 | 0.12 | 0.54 |
| Root Diameter (mm) | Box Cox | 0.0007 | 0.46 | 0.91 | 0.46 | 0.87 | 0.46 |
| Root Length (cm) | Log | 0001.> | 0.90 | 0.08 | 0.90 | 0.06 | 0.90 |

Table S6. The effects of temperature, grafting, and their interaction on the measured traits in PI414723 accession in comparison to self grafted PI414723. Data were transformed where necessary to meet normality assumptions. A probability value below 0.05 indicates a significant difference based on the Student's-T HSD test.

| Treat | Transformation | Temperature | | Treat | | Temp*Treat | |
|---|---|---|---|---|---|---|---|
| | | Prob > F | $R^2$ | Prob > F | $R^2$ | Prob > F | $R^2$ |
| $A_n$ (µmol $m^{-2}$ $s^{-1}$) | Johnson Su | 0001.> | 0.81 | 0.73 | 0.81 | 0.0008 | 0.81 |
| Rn (µmol $m^{-2}$ $s^{-1}$) | Johnson Su | 0.12 | 0.31 | 0.07 | 0.31 | 0.71 | 0.31 |
| $g_{sw}$ (mol $m^{-2}$ $s^{-1}$) | Johnson Su | 0.01 | 0.40 | 0.47 | 0.40 | 0.19 | 0.40 |
| VcMax (µmol $m^{-2}$ $s^{-1}$) | - | 0001.> | 0.87 | 0.59 | 0.87 | 0.23 | 0.87 |
| Jmax (µmol $m^{-2}$ $s^{-1}$) | - | 0.006 | 0.45 | 0.47 | 0.45 | 0.10 | 0.45 |
| Y(II) (a.u.) | - | 0001.> | 0.78 | 0.98 | 0.78 | 0.24 | 0.78 |
| Root Wt (g) | Box Cox | 0001.> | 0.79 | 0.99 | 0.79 | 0.92 | 0.79 |
| Root Diameter (mm) | Box Cox | 0.0026 | 0.49 | 0.91 | 0.49 | 0.10 | 0.49 |
| Root Length (cm) | Log | 0001.> | 0.87 | 0.13 | 0.87 | 0.15 | 0.87 |

Under Review
